# Combination of the parent analogue of Remdesivir (GS-441524) and Molnupiravir results in a markedly potent antiviral effect in SARS-CoV-2 infected Syrian hamsters

**DOI:** 10.1101/2022.10.13.512054

**Authors:** Rana Abdelnabi, Piet Maes, Steven de Jonghe, Birgit Weynand, Johan Neyts

**Affiliations:** KU Leuven Department of Microbiology, Immunology and Transplantation, Rega Institute for Medical Research, Laboratory of Virology and Chemotherapy, B-3000 Leuven, Belgium; The VirusBank Platform, Gaston Geenslaan, B-3000 Leuven, Belgium; Laboratory of Clinical and Epidemiological Virology, Rega Institute, KU Leuven, Department of Microbiology, Immunology and Transplantation, 3000, Leuven, Belgium; Zoonotic Infectious Diseases Unit, Leuven, Belgium; KU Leuven Department of Imaging and Pathology, Translational Cell and Tissue Research, B-3000 Leuven, Belgium; Division of Translational Cell and Tissue Research; Global Virus Network, GVN

**Keywords:** COVID-19, SARS-CoV-2 VoC, GS-441524, Molunpiravir, Antivirals, BA.5, combination

## Abstract

Remdesivir was the first drug to be approved for the treatment of severe COVID-19; followed by molnupiravir (another prodrug of a nucleoside analogue) and the protease inhibitor nirmatrelvir. Combination of antiviral drugs may result in improved potency and help to avoid or delay the development of resistant variants. We set out to explore the combined antiviral potency of GS-441524 (the parent nucleoside of remdesivir) and molnupiravir against SARS-CoV-2. In SARS-CoV-2 (BA.5) infected A549-Dual™ hACE2-TMPRSS2 cells, the combination resulted in an overall additive antiviral effect with a synergism at certain concentrations. Next, the combined effect was explored in Syrian hamsters infected with SARS-CoV-2 (Beta, B.1.351); treatment was started at the time of infection and continued twice daily for four consecutive days. At 4 day 4 post-infection, GS-441524 (50 mg/kg, oral BID) and molnupiravir (150 mg/kg, oral BID) as monotherapy reduced infectious viral loads by 0.5 and 1.6 log_10_, respectively, compared to the vehicle control. When GS-441524 (50 mg/kg, BID) and molnupiravir (150 mg/kg, BID) were combined, infectious virus was no longer detectable in the lungs of 7 out of 10 of the treated hamsters (4.0 log_10_ reduction) and titers in the other animals were reduced by ~2 log_10_. The combined antiviral activity of molnupiravir which acts by inducing lethal mutagenesis and GS-441524, which acts as a chain termination appears to be highly effective in reducing SARS-CoV-2 replication/infectivity. The unexpected potent antiviral effect of the combination warrants further exploration as a potential treatment for COVID-19.

## Introduction

Since it has been declared as a global pandemic by the World Health Organisation (WHO), the coronavirus disease (COVID-19) has continued its devastating impact on public health. Remdesivir (Veklury, Gilead), a broad-spectrum adenosine analog prodrug initially developed for the treatment of the Ebola virus infections, has been the first drug to recieve FDA approval for use in COVID-19 patients. Despite the limited benefits in hospitalized patients (Ader et al., 2022; Consortium, 2020; Goldman et al., 2020), early administration of remdesivir in non-hospitalized COVID-19 patients reduced the risk of hospitalization or death by 87% compared to placebo (Gottlieb et al., 2021). By the end of 2021, two oral antiviral drugs i.e. Molnupiravir (Lagevrio, Merck) and Nirmatrelvir (in combination with ritonavir, Paxlovid, Pfizer) have been authorized by FDA and other countries/regions for emergency use in COVID-19 patients at high risk of hospitalization (Hammond et al., 2022; Syed, 2022).

The parent nucleoside of remdesivir, GS-441524, has been found to be the major circulating metabolite after administration of remdesivir (Warren et al., 2016; Williamson et al., 2020; V. C. Yan & Muller, 2020). GS-441524 exerts antiviral activity (i) in cats infected with the feline coronavirus and (ii) against SARS-CoV-2 infection in AAV-hACE2-transduced mice (Pedersen et al., 2019; Li et al., 2021). It was well-tolerated in cats at doses exceeding that of remdesivir in humans, and in one human subject (Pedersen et al., 2019; V. Yan, 2021). GS-441524 has also been shown to have a favorable oral bioavailability in mice compared to remdesivir (Xie & Wang, 2021).

Molnupiravir (EIDD-2801, MK-4482) is the orally bioavailable counterpart of the ribonucleoside analogue N4-hydroxycytidine (NHC, EIDD-1931), which was initially developed for the treatment of infections with the influenza virus (Toots et al., 2019). NHC exerts *in vitro* antiviral activity against multiple RNA viruses of different families by incorporation into viral RNA, resulting in the accumulation of deleterious mutations in the nascent viral RNA, and consequently, error catastrophe of the viral genome (Urakova et al., 2017). Molnupiravir exerts antiviral activity in SARS-CoV-2 infected mice, Syrian hamsters and ferrets (Cox et al., 2020; Rosenke et al., 2021; Wahl et al., 2020), including against several VoCs (Abdelnabi et al., 2021). Recent data analysis from MOVe-OUT trial (NCT04575597) with molnupiravir showed a relative risk reduction of hospitalization and respiratory interventions in molnpiravir-treated patients of ~34% (Johnson et al., 2022).

Combinations of antiviral drugs, such as against SARS-CoV-2, may allow to achieve a more potent effect than with monotherapy. This amy allow for dose-reductions and result as well as in a reduced risk of resistance development. We previously reported that favipiravir significantly potentiates the antiviral efficacy of molnupiravir when co-adminstered to SARS-CoV-2 infected Syrian hamsters (Abdelnabi et al., 2021a). Similarly, the combined treatment of SARS-CoV-2-infected Syrian hamsters with favipiravir and GS-441524 has been reported to be more efficient as compared to monotherapy with either drug (Chiba et al., 2022). Recently, it has also been reported that combination of suboptimal concentrations of molnupiravir and remdesivir compeletely abrogated the production of SARS-CoV-2 infectious virus particles in the apical wash of human nasal epithelium cultures, although no significant, antiviral effect was detected at the RNA level (Jonsdottir et al., 2022).

Since we previously demonstrated that the combination of molnupiravir and favipiravir results in SARS-CoV-2 infected hamsters in an unexpected potent antiviral effect, we here wanted to assess the combined antiviral potency of the parent nucleoside of remdesivir (GS-441524) with molnupiravir against SARS-CoV-2 infection (*in vitro* and in the hamster infection model).

## Material and methods

### SARS-CoV-2

The SARS-CoV-2 strains used in this study, SARS-CoV-2 Beta B.1.351 variant (derived from hCoV-19/Belgium/rega-1920/2021; EPI_ISL_896474, 2021-01-11) (Abdelnabi, Boudewijns, et al., 2021) and SARS-CoV-2 omicron BA.5 variant [BA.5.2.1, EPI_ISL_14782497] were recovered from nasopharyngeal swabs taken from RT-qPCR confirmed patients. Infectious viruses were first isolated by passaging on Vero E6 cells then passage two virus stocks of the beta variant and the BA.5 variant on Vero E6 and Calu-3 cells, respectively, were used for the studies described here. Live virus-related work was conducted in the high-containment A3 and BSL3+ facilities of the KU Leuven Rega Institute (3CAPS) under licenses AMV 30112018 SBB 219 2018 0892 and AMV 23102017 SBB 219 20170589 according to institutional guidelines.

### Cells

Vero E6 cells (African green monkey kidney, ATCC CRL-1586) were cultured in minimal essential medium (MEM, Gibco) supplemented with 10% fetal bovine serum (Integro), 1% non-essential amino acids (NEAA, Gibco), 1% L-glutamine (Gibco) and 1% bicarbonate (Gibco). A549-Dual™ hACE2-TMPRSS2 cells obtained by Invitrogen (Cat. a549d-cov2r) were cultured in DMEM 10% FCS (Hyclone) supplemented with 10 μg/ml blasticidin (Invivigen, ant-bl-05), 100 μg/ml hygromycin (Invivogen, ant-hg-1), 0.5 μg/ml puromycin (Invivogen, ant-pr-1) and 100 μg/ml zeocin (Invivogen, ant-zn-05). End-point titrations and antiviral assays were performed with media containing 2% fetal bovine serum instead of 10% and no antibiotics. Both cells were kept in a humidified 5% CO_2_ incubator at 37°C.

### Compounds

Molnupiravir (EIDD-2801) and GS-441524 were purchased from Excenen Pharmatech Co., Ltd (China). For *in vitro* assays, 10 mM stocks of compounds were made by dissolving in DMSO. For *in vivo* studies, molnupiravir was formulated as 50 mg/ml stock in a vehicle containing 10% PEG400 (Sigma) and 2.5% Kolliphor-EL (Sigma) in water. GS-441524 was formulated as a 15 mg/mL stock in 30% PEG-400 (Sigma) in PBS containing 1%DMSO.

### *In vitro* evaluation of combined antiviral activity

Combination study was performed by treating A549-Dual™ hACE2-TMPRSS2 cells (15,000 cells/well) with a matrix of 2-fold dilution series of EIDD-1931 and GS-441524 after which the cells were infected with the BA.5 variant at MOI of 0.001. After 4 days of incubation, cell viability was quantified by the MTS/PMS method. Data were then analyzed with the SynergyFinder webtool (Ianevski et al., 2017) based on zero interaction potency (ZIP) model.

### SARS-CoV-2 infection model in hamsters

The hamster infection model of SARS-CoV-2 has been described before (Abdelnabi et al., 2022). Female Syrian hamsters (Mesocricetus auratus) were purchased from Janvier Laboratories and kept per two in individually ventilated isolator cages (IsoCage N Bio-containment System, Tecniplast) at 21°C, 55% humidity and 12:12 day/night cycles. Housing conditions and experimental procedures were approved by the ethics committee of animal experimentation of KU Leuven (license P065-2020). For infection, female hamsters of 6-8 weeks old were anesthetized with ketamine/xylazine/atropine and inoculated intranasally with 50 μL containing 1×10^4^ TCID50 SARS-CoV-2 beta variant (day 0). On day 4 post-infection, animals were euthanized for sampling of the lungs and further analysis by intraperitoneal (i.p.) injection of 500 μL Dolethal (200 mg/mL sodium pentobarbital).

### *In vivo* treatment regimen

Hamsters were treated twice daily (BID) via oral gavage with either vehicle, 150 mg/kg EIDD-2801, 50 mg/kg GS-441524 or combination of both compounds starting from the time of infection (d0) with SARS-CoV-2 beta variant. All the treatments continued until day 3 post-infection (pi). Hamsters were monitored for appearance, behavior and weight. At day 4 pi, hamsters were euthanized as mentioned earlier. Lungs were collected for viral loads determination and histopathology scoring. Viral RNA and infectious virus were quantified by RT-qPCR and end-point virus titration, respectively.

### SARS-CoV-2 RT-qPCR

Hamster lung tissues were collected after sacrifice and were homogenized using bead disruption (Precellys) in 350 μL TRK lysis buffer (E.Z.N.A.® Total RNA Kit, Omega Bio-tek) and centrifuged (10.000 rpm, 5 min) to pellet the cell debris. RNA was extracted according to the manufacturer’s instructions. RT-qPCR was performed on a LightCycler96 platform (Roche) using the iTaq Universal Probes One-Step RT-qPCR kit (BioRad) with N2 primers and probes targeting the nucleocapsid (Boudewijns et al., 2020). Standards of SARS-CoV-2 cDNA (IDT) were used to express viral genome copies per mg tissue (Kaptein et al., 2020).

### End-point virus titrations

Lung tissues were homogenized using bead disruption (Precellys) in 350 μL minimal essential medium and centrifuged (10,000 rpm, 5min, 4°C) to pellet the cell debris. To quantify infectious SARS-CoV-2 particles, endpoint titrations were performed on confluent Vero E6 cells in 96-well plates. Viral titers were calculated by the Reed and Muench method (Reed & Muench, 1938) using the Lindenbach calculator and were expressed as 50% tissue culture infectious dose (TCID50) per mg tissue.

### Histology

For histological examination, the lungs were fixed overnight in 4% formaldehyde and embedded in paraffin. Tissue sections (5 μm) were analyzed after staining with hematoxylin and eosin and scored blindly for lung damage by an expert pathologist. The scored parameters, to which a cumulative score of 1 to 3 was attributed, were the following: congestion, intra-alveolar hemorrhagic, apoptotic bodies in bronchus wall, necrotizing bronchiolitis, perivascular edema, bronchopneumonia, perivascular inflammation, peribronchial inflammation and vasculitis.

### Statistics

The detailed statistical comparisons, the number of animals and independent experiments that were performed is indicated in the legends to figures. “Independednt experiments” means that experiments were repeated separately on different days. The analysis of histopathology was done blindly. All statistical analyses were performed using GraphPad Prism 9 software (GraphPad, San Diego, CA, USA). Statistical significance was determined using the non-parametric Mann Whitney U-test. P-values of <0.05 were considered significant.

### Ethics

Housing conditions and experimental procedures were done with the approval and under the guidelines of the ethics committee of animal experimentation of KU Leuven (license P065-2020).

### Data Availability

All of the data generated or analyzed during this study are included in this published article.

## Results

First, the dose-response effects of the active metabolite of molnupiravir (EIDD-1931) and GS-441524 were determined against the BA.5 variant in A549-Dual™ hACE2-TMPRSS2 cells using a cytopathic effect (CPE) reduction asssay. Both EIDD-1931 (Fig. 1A) and GS-441524 (Fig. 1B) efficiently inhibited the BA.5 variant replication with EC50 values of 0.96 ± 0.13 and 2.1 ± 0.27 μM, respectively.

**Fig. 1.**
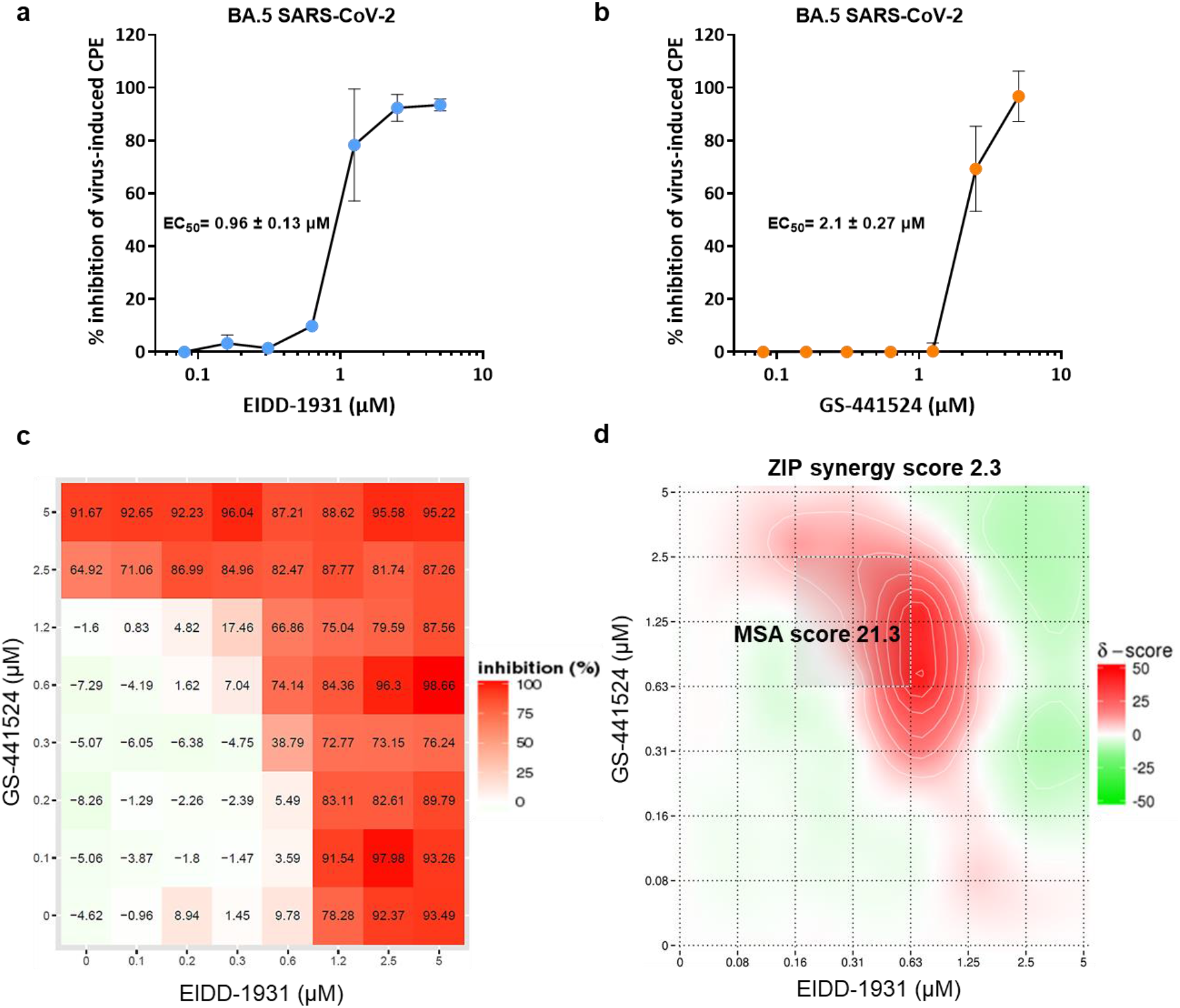
*In vitro* efficacy of the combination of EIDD-1931 and GS-441524 against SARS-CoV-2 BA.5 variant. Dose-response Dose-response effect of (A) EIDD-1931 and (B) GS-441524 on SAR-CoV-2 (BA.5 variant)-induced cytopathic effect (CPE) at MOI 0.001, as quantified in A549-Dual™ hACE2-TMPRSS2 cells by the MTS/PMS method. Data represented in tehj graphs are mean values ± standard deviations (SD) from two independent experiments. (C) Dose-response matrix for EIDD-1931 and GS-441524 representing %inhibition of virus-induced CPE in BA.5-infected A549-Dual™ hACE2-TMPRSS2 cells. (D) heat map of the delta scores (%) for the combined EIDD-1931 and GS-441524 antiviral effects based on ZIP analysis where δ = 0, δ > 0, and δ < 0 correspond to zero interaction, synergy, and antagonism, respectively. The overall ZIP score represents the response beyond expectation (in %). In the range −10 < ZIP < 10, the compounds are likely to act in an additive manner, Score ≥10 indicate syenrgism. MSA= most synergistic area. The graphics for combination represent the means of two independent experiments.

Next, the combined antiviral activity of EIDD-1931 and GS-441524 against the BA.5 variant was assessed in a checkerboard (matrix) format with seven serial dilutions of each compound. The inhibition of virus-induced CPE by each compound alone or in combination was determined (Fig. 1C), and the data were analyzed with the SynergyFinder webtool (Ianevski et al., 2017) based on zero interaction potency (ZIP) model (Fig. 1D). The EIDD-1931 and GS-441524 combination resulted in an overall synergy score of 2.28 (Fig. 1D). A large surface of the combined antiviral effect of both compound resulted in additive antiviral effects (Fig. 1D). However, at certain concentration ranges, the combinations resulted in a marked synergistic effect with a score of 21.32 in the most synergistic area (Fig. 1D). No cytotoxicity was observed for at any combined concentrations of both compounds.

Next, we evaluated the efficacy of single versus combination treatment with molnupiravir [EIDD-2801] (150 mg/kg, oral BID) and GS-441524 (50 mg/kg, oral BID) in SARS-CoV-2-infected hamsters. Briefly, hamsters were treated with vehicle or the intended dose of each compound as single or combined therapy for four consecutive days starting just before the infection with SARS-CoV-2 beta (B.1.351) variant (Fig. 2A). At day 4 post-infection (pi), the animals were euthanized and organs were collected for quantification of viral RNA, infectious virus titers and to assess the impact on lung histopathology (Fig. 2A). Single treatment with molnupiravir (150 mg/kg BID) significantly reduced viral RNA and infectious virus titers in the lungs by 1.3 (P=0.0045) and 1.6 (P=0.0003) log_10_/mg lung tissue, respectively (Fig. 2B/C). As monotherapy, GS-441524 treatment at dose of 50 mg/kg (BID) reduced viral RNA and infectious virus titers in the lung by 1.2 (P=0.0001) and 0.5 (P=0.043) log_10_/mg lung tissue, respectively, compared to the vehicle-treated hamsters (Fig. 2B/C). On the other hand, the combined molnupiravir/GS-441524 treatment resulted in more significant reduction of of lung viral RNA loads [2.4 log_10_ (P<0.0001)] compared to single treatment with molnupiravir (150 mg/kg, BID) (Fig. 2B). Interestingly, the molnupiravir/GS-441524 combination resulted in a markedly enhanced reduction in infectious virus titers (~4.0 log_10_ TCID50 per mg lung, P<0.0001 as compared to molnupiravir alone) (Fig. 2C). Notably, there was no detectable infectious virus in the lungs of 7 out of 10 hamsters in the combined treatment group (Fig. 2C). A significant improvement in the cumulative histological lung pathology scores was observed in the combined treatment molnupiravir/GS-441524 group (median histopathology score of 2.5) as compared to the vehicle control (median score of 4.5), P=0.0002 (Fig. 2D). The improvement in lung histopathology in the combined molnupiravir/GS-441524 treatment group was also significant compared to the single molnupiravir treatment group (median score of 3.5, P=0.04) as well as better than the single GS-441524 treatment group (median score of 3, P=0.13, non-significant) (Fig. 2D). No significant weight loss or toxicity signs were observed in either the single treatment or the molnupiravir/GS-441524 combination groups compared to the vehicle-treated groups (Fig. 2E). Animals treated with the combination therapy had the highest, although not significant, weight gain (mean % weight change of 4.5), compared to the other groups (Fig. 2E).

**Fig. 2.**
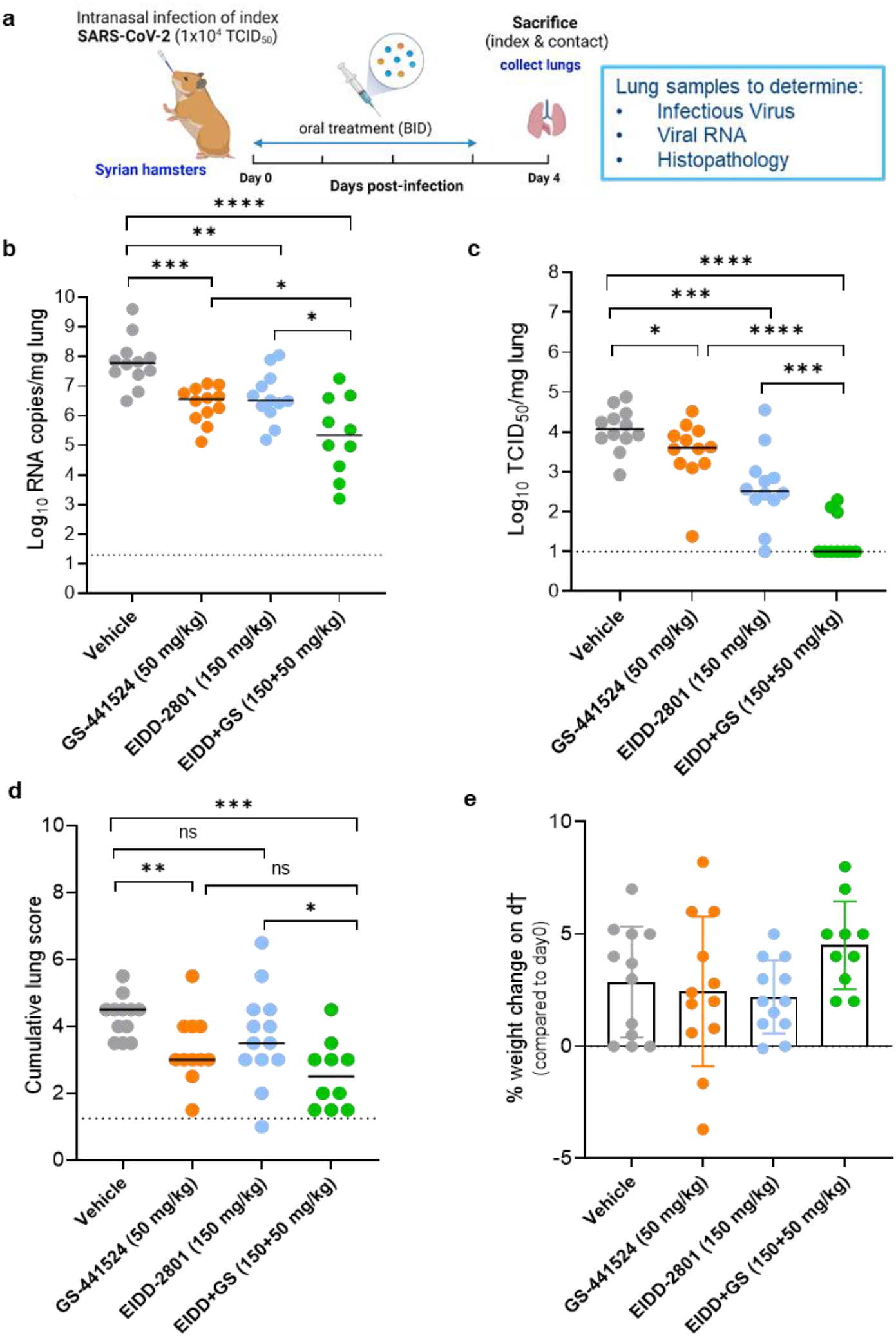
Combined efficacy of Molnupiravir (EIDD-2801) with GS-441524 against SARS-CoV-2 B.1.351 variant in a hamster infection model. (A) Set-up of the study. (B) Viral RNA levels in the lungs of control (vehicle-treated), EIDD-2801-treated (150 mg/kg, BID), GS-441524-treated (50 mg/Kg, BID) and combination-treated (EIDD-2801+GS-441524 at 150+50 mg/kg BID) SARS-CoV-2 (B.1.351)-infected hamsters at day 4 post-infection (pi) are expressed as log_10_ SARS-CoV-2 RNA copies per mg lung tissue. Individual data and median values are presented. (C) Infectious viral loads in the lungs of control (vehicle-treated), EIDD-2801-treated, GS-441524-treated and combination-treated (EIDD-2801+ GS-441524) SARS-CoV-2-infected hamsters at day 4 pi are expressed as log_10_ TCID50 per mg lung tissue. Individual data and median values are presented. (D) Cumulative severity score from H&E stained slides of lungs from different treatment groups. Individual data and median values are presented and the dotted line represents the median score of untreated non-infected hamsters. (E) Weight change at day 4 pi in percentage, normalized to the body weight at the time of infection. Bars represent means ± SD. Data were analyzed with the Mann-Whitney U test. *P < 0.05, **P < 0.01, ***P < 0.001, ****P < 0.0001, ns=non-significant, i.e. P<0.05). EIDD=EIDD-2801. All data (panels B, C, D, E) are from two independent experiments with 12 animals per group except for the combination group (n=10). Panel A created by Bioredner.

## Discussion

Combinations of antiviral drugs proved to be more efficient than monotherapy for the treatment of viral infections such as caused by the human immunodeficiency virus (HIV) and the hepatitis C virus (HCV). It allows also to prevent the emergence of drug-resistant variants (Shyr et al., 2021). Here, we explored the potential efficacy of the combined GS-441524 and molnupiravir treatment against SARS-CoV-2 variants of concerns both *in vitro* and in infected hamsters.

In SARS-CoV-2 (BA.5) infected A549-Dual™ hACE2-TMPRSS2 cells, the combination resulted in an overall additive antiviral effect with a marked synergism at certain concentrations. In hamsters infected with the beta (B.1.351) variant and that received suboptimal doses of either GS-441524 or molnupiravir, limited to moderate antiviral effects (measured as reduction of viral titers in the lungs) were noted, respectively. However, the combined treatment of both compounds resulted in a potent antiviral effect with low or in most animals undetectable titers of infectious virus in the lungs.

Molnupiravir has been shown to increase the mutation frequency of MERS-CoV viral RNA in infected mice (Sheahan et al., 2020); we made similar observations in SARS-CoV-2-infected hamsters (Abdelnabi, et al., 2021b). These data strongly indicate that molnupiravir exerts its antiviral activity by inducing an error catastrophe of the replicating SARS-CoV-2 genome. Similar observations were made in studies in mice infected with the Venezuelan equine encephalitis virus or the influenza virus (Jordan et al., 2018; Urakova et al., 2017). GS-441524 (the active substance of remdesivir), is metabolized in cells into the pharmacologically active nucleoside triphosphate, which stalls viral RNA synthesis after being incorporated into the growing RNA strand by the RdRp (Kokic et al., 2021). Therefore, the combined actions of inducing lethal mutagenesis (molnupiravir) in the virus with chain termination (GS-441524) appears to be highly effective in reducing SARS-CoV-2 replication and in particular, infectivity.

To conclude, we identified a potent and well-tolerated combination of the nucleoside analogues, molnupiravir and GS-441524 against SARS-CoV-2. Carefully designed combinations may result in potent antiviral effects and (although not studied here) may also reduce the risk of drug-resistance development. This study lays the basis for further exploration of the combined treatment of SARS-CoV-2 infections with molnupiravir and new oral prodrug versions of remdesivir such as GS-621763 (Schäfer et al., 2022).

## Acknowledgments

We thank Carolien De Keyzer, Lindsey Bervoets, Thibault Francken, Birgit Voeten, Stijn Hendrickx, Niels Cremers for excellent technical assistance. We are grateful to Piet Maes for kindly providing the SARS-CoV-2 variants used in this study. We thank Prof. Jef Arnout and Dr. Annelies Sterckx (KU Leuven Faculty of Medicine, Biomedical Sciences Group Management) and Animalia and Biosafety Departments of KU Leuven for facilitating the animal studies.

## Author Contributions

R. A. and J.N. designed the studies; R.A and B.W. performed the studies and analyzed data; R.A. made the graphs; B.W. and J.N. provided advice on the interpretation of data; R.A. and J.N. wrote the paper; S. D. Jand P.M. provided essential reagents; R.A. and J.N. supervised the study; J.N. acquired funding.

